# Fiber-Coupled Rotating Two-Photon Light-Sheet Microscopy for Physiological Vestibular Imaging in Larval Zebrafish

**DOI:** 10.64898/2026.10.10.756414

**Authors:** Hugo Trentesaux, Geoffrey Migault, Thomas Panier, Georges Debregeas, Volker Bormuth

**Author notes:** These authors contributed equally to this work (Hugo Trentesaux, Geoffrey Migault, Thomas Panier).

## Abstract

Physiological vestibular stimulation requires macroscopic rotation of the animal, making simultaneous high-resolution neural imaging technically demanding. Here we present a fiber-coupled rotating two-photon light-sheet microscope for physiological pitch-axis vestibular imaging in larval zebrafish. Two-photon excitation is attractive in this context because near-infrared excitation avoids visible-light contamination of the visual environment. Femtosecond pulses were delivered to the rotating microscope through a broadband hollow-core negative-curvature fiber previously characterized in static light-sheet configurations. The rotating instrument maintained the axial mechanical stability required for cellular-resolution imaging, with submicron mean axial displacement over the characterized angular range. Fiber performance under rotation was also stable: bend-induced transmission changes were small in the relevant curvature regime, and circular input polarization suppressed rotation-dependent polarization artefacts to below 1% in the sample plane. The dominant rotation-sensitive optical limitation was not the fiber but perturbation of the thermal lens in the sample chamber. Reducing fluid motion along the excitation path suppressed this artefact and restored stable imaging during dynamic rotation. With this system, we recorded brain-wide vestibular responses during sinusoidal pitch stimulation and resolved anatomically organized cellular response patterns across the brain. These results establish rotating two-photon light-sheet microscopy as a practical solution for vestibular imaging in experiments that require physiological stimulation together with darkness or controlled visual conditions.

## 1. Introduction

The vestibular system reports self-motion and head orientation and contributes critically to gaze stabilization, posture, balance, and spatial perception [1, 2]. Because vestibular processing engages distributed sensorimotor circuits, addressing it benefits from brain-scale, ideally whole-brain, neural recordings. Larval zebrafish are particularly well suited to this problem because light-sheet microscopy enables near-cellular-resolution functional imaging across large fractions of the brain *in vivo* [3, 17–19].

Several groups have used this preparation to study distributed neural activity under controlled sensory conditions, including visual stimulation and fictive behavior [3, 17, 19]. For vestibular experiments, however, the motion required for physiological stimulation directly perturbs conventional imaging approaches [14]. Stationary alternatives such as optical trapping or magnetic actuation of the otoliths are powerful because they provide experimentally tractable substitutes for vestibular drive in head-fixed animals, but they reproduce inertial stimulation indirectly and with different calibration constraints [4–6].

We previously developed a miniaturized rotating one-photon light-sheet microscope that delivered physiological pitch and roll stimulation while preserving whole-brain imaging stability [14, 25]. Other microscope configurations, that provide physiological vestibular stimulations are tiltable-objective microscopy [7] and tilt-in-place microscopy [8]. Although one-photon imaging successfully records vestibular responses, its visible excitation compromises sensitivity to additional visual stimuli: a same-fish comparison demonstrated attenuated flash-evoked responses under 488 nm relative to 930 nm light-sheet excitation [18]. Two-photon excitation therefore offers an advantage when the aim is to measure responses to visual stimuli alongside brain-wide vestibular activity.

Two-photon light-sheet microscopy addresses this limitation by moving the excitation wavelength outside the visual range of the animal while retaining optical sectioning for brain-scale imaging [18]. The main obstacle in a rotating implementation is then the reliable delivery and shaping of femtosecond pulses on a moving microscope platform. Negative-curvature hollow-core fibers are attractive for this task because they provide broadband transmission of ultrashort pulses with low dispersion, low nonlinearity, and good mode quality [9, 10, 15]. Conventional solid-core fibers are much less suitable for femtosecond delivery because they introduce stronger dispersion, nonlinear effects, and polarization- and bending-dependent fluctuations [9]. Using this fiber technology, we recently presented a compact fiber-coupled one- and two-photon light-sheet microscope in a static configuration [15], but did not address the specific constraints imposed by dynamic rotation. In a rotating vestibular microscope, this problem is especially stringent because any rotation-coupled change in excitation power, polarization state, or beam position is synchronized to the stimulus waveform and can therefore mimic neural activity. Whether these effects can be suppressed sufficiently for stable brain-wide two-photon vestibular imaging remained unresolved.

Here we present a rotating two-photon light-sheet microscope for physiological pitch-axis vestibular stimulation in larval zebrafish. We first describe the compact optical and mechanical design and show that the rotating microscope provides the mechanical stability required for cellular-resolution imaging. We then quantify fiber behavior under rotation and find that bend-induced transmission and polarization changes are negligible under the operating conditions of the instrument. Next, we identify rotation-dependent perturbation of the thermal lens in the sample chamber as the dominant optical limitation and show that shortening the laser propagation path through water and constraining fluid motion suppresses this artefact for functional imaging. Finally, we use the stabilized system for whole-brain two-photon imaging during sinusoidal pitch stimulation and recover anatomically organized vestibular response maps consistent with previous rotating one-photon measurements [14, 25].

## 2. Results

### 2.1. Rotating two-photon light-sheet microscope design and imaging stability

The rotating microscope was built on the compact mechanical architecture of our previous one-photon vestibular microscope [14], but with a redesigned optical layout for two-photon operation and a different actuation mechanism. In the present implementation, the light-sheet unit was mounted on the rear side of the rotating breadboard, an additional mirror was introduced to align the fiber outlet with the mechanical rotation axis, and pitch stimulation was generated by a continuously driven motor via a linkage mechanism that converted continuous motor rotation into sinusoidal pitch motion (Fig. 1A,B). A femtosecond laser at 915 nm for two-photon excitation was delivered to the rotating platform through a broadband hollow-core negative-curvature fiber [15], and the optical assembly was aligned with the mechanical rotation axis for pitch stimulation (Fig. 1A,B). Technical implementation details are provided in Methods. The static optical performance of the two-photon light-sheet design on which the present system is based was characterized previously [15], including the light-sheet profile measured with submicron fluorescent beads. The measured light-sheet thickness was 4.5 µm, corresponding to a two-photon excitation width of approximately 3 µm.

**Fig. 1.**
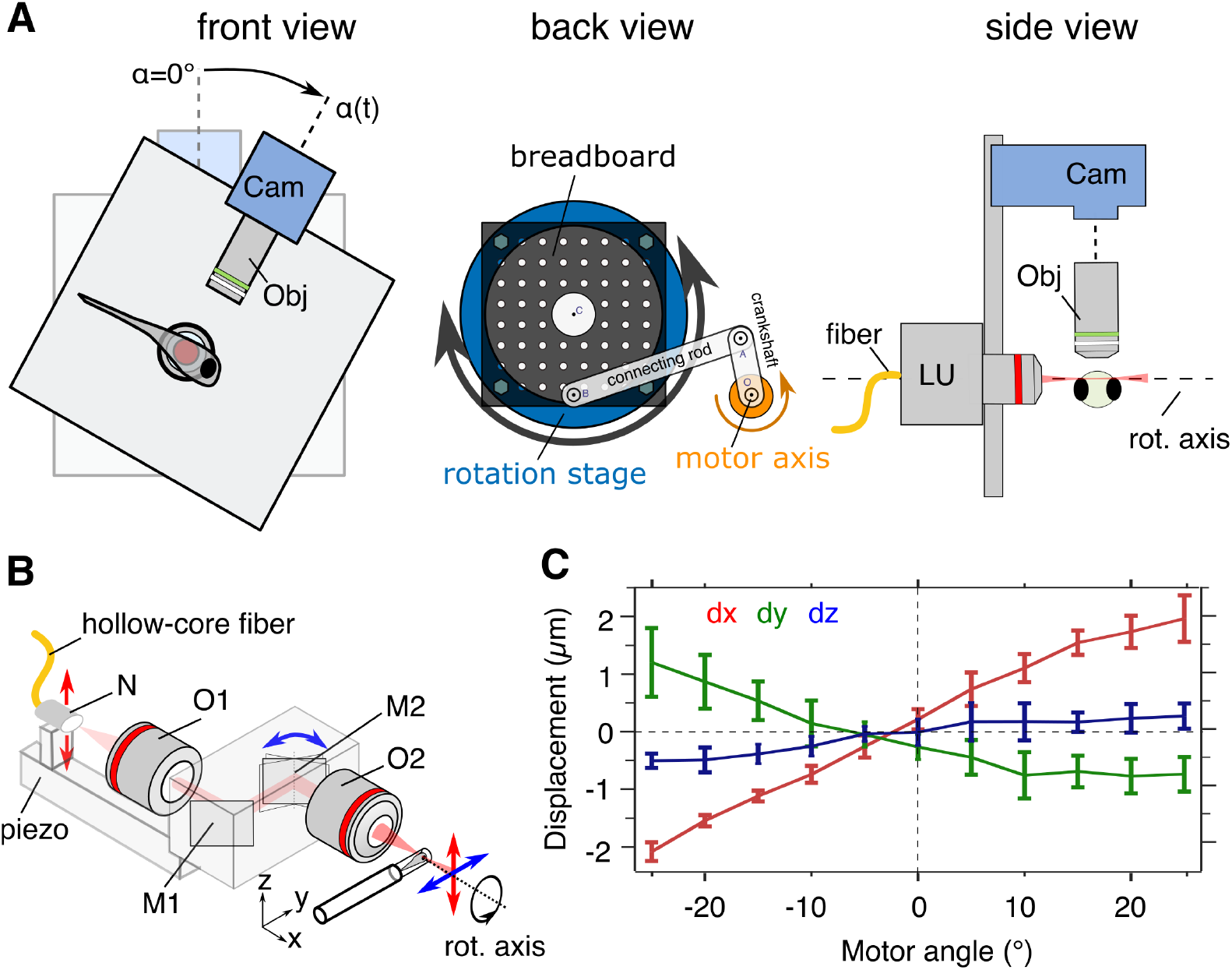
Rotating two-photon light-sheet microscope and mechanical stability during pitch-axis vestibular stimulation. (A) Front, back, and side views of the rotating microscope showing the orientation of the larval zebrafish with respect to the pitch axis, the compact optical arrangement, and the rear-side motor-driven pitch-actuation mechanism. (B) Schematic of the compact two-photon light-sheet unit mounted on the rotating breadboard and aligned with the mechanical rotation axis. Abbreviations: N, NA expander; O1, collimation objective; M1, fixed mirror; M2, scanning mirror; O2, illumination objective. (C) Relative displacement of a microscopic bead placed at the sample position as a function of microscope angle, showing mean displacement and standard deviation in *x* (red), *y* (green), and *z* (blue). The mean axial displacement (*z*) remains submicron, while the peak-to-peak lateral excursions are approximately 4 µm in *x* and 2 µm in *y* over the characterized angular range of ±25°.

We quantified mechanical stability directly at the sample position using microscopic beads, because any deformation of the mechanical structure during rotation would produce a relative displacement between the excitation and detection planes and the sample, which would be detrimental for cellular-resolution functional imaging. Over the characterized angular range of ±25°, mean displacement ranged approximately from −2 to +2 µm in *x* and from −0.8 to +1.2 µm in *y*, while remaining within approximately 0.5 µm of the reference position along the detection axis (*z*; Fig. 1C). Axial stability is particularly important because out-of-plane motion changes the sampled tissue and cannot be corrected by lateral image registration. Lateral displacement was corrected during post-processing using the procedure described previously [14]. Functional imaging was performed over the smaller angular range of ±10°.

### 2.2. Fiber bending and polarization effects are negligible in the operating regime

The hollow-core fiber used here allows broadband delivery of ultrashort pulses with low dispersion and low attenuation [15]. In the rotating microscope, however, platform motion bends and twists the fiber in synchrony with the vestibular stimulus. Any resulting change in transmitted power or output polarization would therefore be temporally locked to the stimulation waveform and could be misinterpreted as stimulus-evoked fluorescence changes.

We first quantified bend-dependent transmission losses as a function of imposed curvature using the measurement scheme shown in Fig. 2A. The measured attenuation increased approximately with the square of the fiber curvature, in agreement with analytical expectations for hollow-core guidance [20, 21] (Fig. 2B). Representative output mode profiles across the measured curvature range are shown in Fig. 2B, indicating stable near-single-mode guidance under these conditions. We then addressed polarization sensitivity. In light-sheet microscopy, detected fluorescence can depend on the polarization state of the excitation beam because the illumination path is orthogonal to the detection path. Our fiber is not polarization-maintaining, and the orientation of the output polarization changes with bending. Additional curvature-dependent polarization and ellipticity measurements are shown in Supplementary Fig. S1. To suppress this source of artefact, we coupled circularly polarized light into the fiber and quantified the residual power fluctuations during realistic microscope rotation both with and without a linear polarizer at the output (Fig. 2C,D). The measured power modulation remained below 1% during sinusoidal rotation (Fig. 2D).

**Fig. 2.**
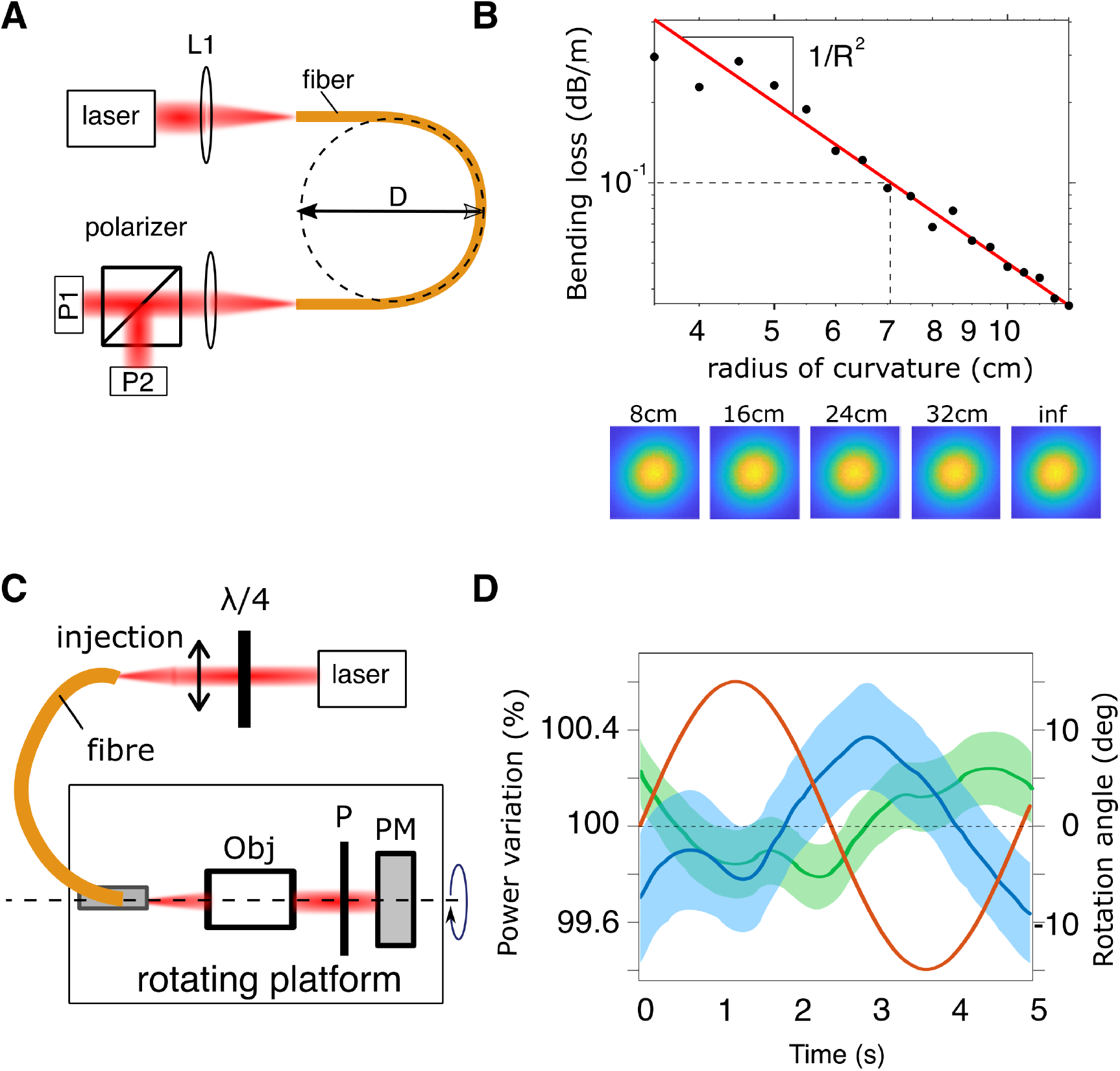
Hollow-core fiber performance under bending and rotation. (A) Bend-loss assay used to quantify transmitted power as a function of fiber curvature. L1, coupling lens; P1 and P2, power meters. (B) Bend-dependent transmission loss of the hollow-core negative-curvature fiber. Loss is plotted in dB/m and scales approximately with the square of fiber curvature. Representative output mode profiles indicate stable near-single-mode guidance over the measured curvature range. (C) Rotation-coupled measurement scheme used to quantify power stability at the output of the rotating light-sheet module after circular input polarization and output polarization analysis. (D) Relative transmitted power (left axis) during sinusoidal rotation, measured with (blue) or without (green) an output polarizer (P) before the power meter (PM). Shaded bands indicate ±1 standard deviation around the mean. The red curve shows the rotation angle (right axis). Power modulation remains below 1% with circular input polarization.

### 2.3. Rotation-dependent thermal lensing in the sample chamber is the dominant optical limitation

Despite the mechanical stability of the microscope and the small fiber-induced fluctuations described above, initial whole-brain calcium recordings during sinusoidal pitch rotation showed widespread fluorescence modulation locked to the stimulus frequency. Phase maps, which represent the phase of fluorescence modulation at the stimulation frequency relative to the stimulus (Methods), revealed broad regions of differing phase across the two brain hemispheres that overshadowed anatomically localized cellular responses (Fig. 3A), suggesting a large-scale illumination artefact. The artefact was negligible at very slow stimulation (< 0.1 Hz) but increased with rotation frequency and acceleration.

**Fig. 3.**
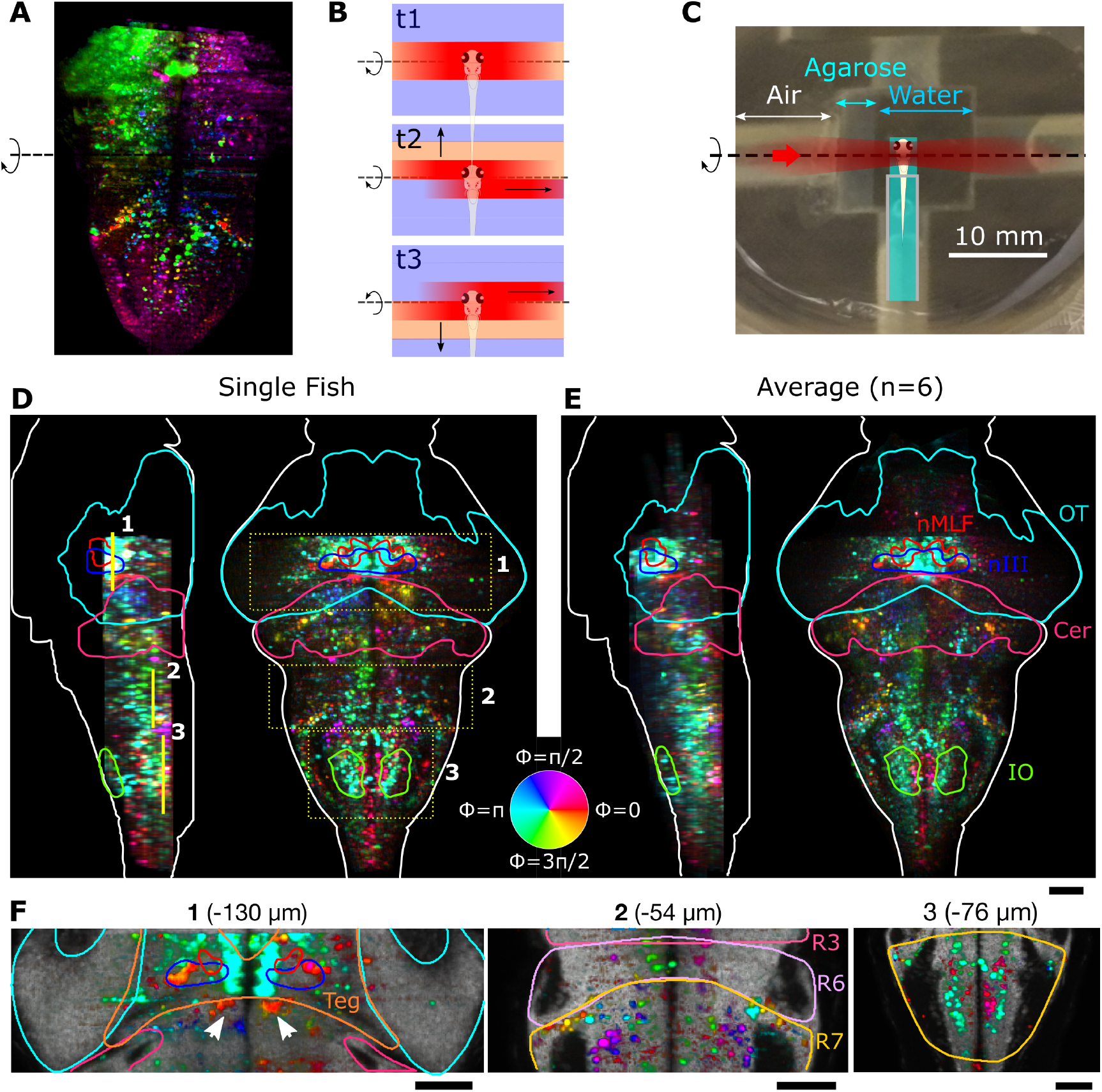
Rotation-dependent thermal lensing and whole-brain vestibular imaging in two-photon mode. (A) Example phase map illustrating the artefact observed during rotation before thermal stabilization of the sample chamber. (B) Schematic interpretation of the rotation-dependent optical instability in the sample chamber. Rotation perturbs the temperature profile around the excitation path and thereby displaces the light-sheet waist. (C) Reduced sample chamber, approximately 10 mm on a side, with added agarose between the laser entry window and the fish to limit water motion along the excitation path. (D) Example phase maps from a single larva during sinusoidal pitch stimulation imaged in two-photon mode, shown in orthogonal views. (E) Average phase maps across six larvae after registration to a common anatomical reference. Labeled regions include the oculomotor nucleus (nIII), nucleus of the medial longitudinal fasciculus (nMLF), optic tectum (OT), cerebellum (Cer), and inferior olive (IO). (F) Enlarged views from representative regions of interest highlighted in (D), showing single-cell-resolved vestibular response patterns. Scale bars are 50 µm. Visualization 1 is related to panel (B) and visualization 2 is related to panel (D).

To identify the origin of this effect, we imaged the light-sheet in fluorescein solution during rotation. We observed a perturbation front traveling across the illuminated plane together with a transient displacement of the beam waist along the propagation axis (see Visualization 1, related to Fig. 3B). From these observations, we identified a thermal lens effect in the sample chamber as the likely origin of the artefact. We subsequently quantified power- and water-path-dependent focal shifts and thermal transients in a dedicated experimental and theoretical study [16]. In brief, absorption of the excitation beam heats the water along the optical path and creates a temperature-dependent refractive-index gradient that acts as a weak diverging lens. In that conduction-based model, the steady-state focal shift is approximated as

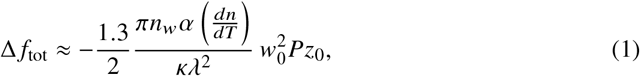

where *n*_*w*_ is the refractive index of water, α its absorption coefficient at the excitation wavelength, *dn*/*dT* the temperature dependence of the refractive index, *κ* the thermal conductivity of water, λ the excitation wavelength, *w*_0_ the beam waist radius, *P* the laser power, and *z*_0_ the length of water traversed by the beam before the focus.

This expression shows that the thermal-lens-induced focal shift increases with absorbed power and with the water-path length *z*_0_. The temporal response of the thermal lens is likewise spatially heterogeneous because the local equilibration time τ increases with the square of the local beam radius. Near the waist, where the beam radius is smallest, equilibration therefore occurs rapidly within milliseconds, whereas more distant water layers along the excitation path, especially near the chamber entrance where the beam is widest, respond on much longer timescales within seconds. In the rotating microscope, these timescales suggest that slow rotation allows the temperature profile to remain close to equilibrium, whereas faster rotation can perturb it before it has fully re-equilibrated, consistent with the observed increase in artefact amplitude. Reducing *z*_0_ shortens the water path ahead of the focus and thus lowers the thermal-lens-induced focal shift while also removing the most slowly responding regions from the illuminated volume, thereby accelerating the overall thermal-lens dynamics.

Accordingly, we reduced the sample-chamber dimensions to approximately 10 mm on a side, thereby reducing the propagation distance through water before the excitation beam reaches the fish to approximately 5 mm. We further filled most of this path with agarose from the laser entry window towards the fish, leaving less than 1 mm of free water immediately upstream of the fish in which rotation-induced fluid motion could perturb the thermal profile (Fig. 3C).

### 2.4. Whole-brain two-photon imaging during physiological pitch-axis vestibular stimulation

With the optimized setup, we performed whole-brain functional imaging during sinusoidal pitch stimulation. Head-restrained larval zebrafish were rotated together with the microscope while fluorescence was recorded in two-photon mode. Visualization 2 shows the spatiotemporal evolution of Δ*F*/*F* through one stimulation cycle (averaged over 120 periods), displayed for 12 axial planes covering 110 µm of brain depth in 10 µm steps. Acquisition and stimulation details are given in Methods.

The widespread phase pattern observed before chamber modification (Fig. 3A) was no longer apparent. Instead, the resulting phase maps resolved individual cell bodies and revealed bilaterally organized vestibular response patterns across the brain (Fig. 3D–F). Response maps from individual fish were highly structured, and the average map across six larvae retained clear anatomical organization. Prominent responsive populations were observed in the tegmentum, including regions consistent with the oculomotor nucleus III and the nucleus of the medial longitudinal fasciculus, as well as in the cerebellum, optic tectum, inferior olive, and hindbrain. Representative enlarged views from selected regions further show clearly resolved cellular response patterns in tegmental and hindbrain structures (Fig. 3F). The anatomical distribution agrees qualitatively with previously published rotating one-photon pitch-tilt response maps [25], providing a biological reference for the present recordings.

Together, these results demonstrate that a rotating two-photon light-sheet microscope can support cellular-resolution whole-brain vestibular imaging during physiological pitch stimulation, provided that the sample chamber is configured to suppress rotation-dependent perturbation of the thermal lens.

## 3. Discussion

We show that whole-brain two-photon light-sheet imaging during physiological pitch-axis vestibular stimulation is technically feasible in larval zebrafish. The main methodological conclusion is that, in a rotating two-photon light-sheet microscope, the dominant rotation-sensitive instability can arise in the thermally loaded sample chamber rather than in hollow-core-fiber delivery itself. Under the bending and twisting conditions imposed by the platform, the fiber remained stable enough in transmission and polarization for calcium imaging, whereas rotation-dependent perturbation of the thermal lens in the chamber was the dominant source of artefact. This result shifts the engineering focus from beam delivery alone to the coupled optical and fluidic behavior of the sample chamber during rotation.

This result also helps position the present instrument within the broader set of methods now available for vestibular imaging in larval zebrafish. Stationary fictive-stimulation strategies such as optical trapping or magnetic actuation of the otoliths are in principle well matched to two-photon light-sheet microscopy, because they avoid moving the microscope and sample chamber during imaging [4–6]. In contrast, approaches based on real spatial reorientation of the animal, including tilt-in-place or rotating configurations, are likely to face related optical constraints because motion of the sample and surrounding fluid can perturb the thermal state along the light-sheet excitation path [7, 8, 14].

More broadly, the results define a concrete engineering rule for rotating multiphoton microscopy: beam delivery and sample-chamber optics must be optimized together. The balance between fiber stability and chamber-induced optical instability is geometry- and power-dependent and should therefore be assessed for each new configuration. Importantly, the relevant instability is not only a steady-state problem, but also depends on whether the thermal profile can re-equilibrate on the timescale of the imposed motion.

The static pulse-delivery and mode characterization reported previously [15], together with the bending-dependent mode profiles and rotation-coupled power measurements presented here, support negative-curvature hollow-core fiber as a practical delivery solution for this instrument. Polarization-maintaining or single-polarization hollow-core fiber designs could reduce rotation-coupled polarization sensitivity further at the delivery stage [11, 12]. In addition, as discussed in our earlier static platform paper [15], the broadband transmission of hollow-core delivery is practically useful because it supports coupling of visible guide light for alignment and sample exploration in addition to femtosecond excitation, while preserving flexibility to adapt the same architecture to other excitation wavelengths and optical reporters [13].

The recovery of vestibular response patterns consistent with previously published one-photon recordings [14, 25] validates the instrument for brain-wide two-photon functional imaging during physiological pitch stimulation. This capability provides a basis for future studies combining vestibular and visual stimulation while preserving sensitivity to visual stimuli that would otherwise be compromised by visible light-sheet excitation [18].

## 4. Materials and methods

### 4.1. Animals

All experiments were performed on 5–7 days post-fertilization larval zebrafish maintained at 28 °C on a 14 h/10 h light/dark cycle. Eggs were collected in the morning and raised in E3 medium. Larvae were fed from 5 dpf onward. Functional imaging experiments were performed in the nacre background using a pan-neuronal, nucleus-localized calcium indicator line, Tg(elavl3:H2B-GCaMP6f), kindly provided by M. B. Ahrens and previously described in the context of light-sheet imaging [22]. All procedures were approved by Le Comité d’Éthique pour l’Expérimentation Animale Charles Darwin C2EA-05 (02601.01 and #32423-202107121527185 v3).

### 4.2. Rotating two-photon light-sheet microscope

The microscope was built on a rotating breadboard carrying the light-sheet illumination unit, the detection optics, and the sample. The optical architecture was derived from our previously published rotating vestibular microscope [14] and from our later static one-/two-photon light-sheet unit [15]. In the present implementation, the animal was rotated about the pitch axis.

Excitation was provided by a femtosecond tunable laser [MaiTai, 90077043, Spectra-Physics, USA]. The beam was circularly polarized using a quarter-wave plate [WPQSM05-915, Thorlabs, USA] and coupled into a hollow-core negative-curvature fiber [PMC-C-9005 B2, GLOphotonics, France] using an injection lens [AC254-040-B-ML, Thorlabs, USA]. Fiber coupling and pulse precompensation followed the procedures described in [15], which also reports the fiber’s transmission properties under static conditions.

The light-sheet unit on the rotating platform comprised an NA expander lens [C151TMD-B, Thorlabs, USA], a collimation objective [LMPlan N 5x/0.10 IR, Olympus, Japan], a fixed mirror [BB0511-E03, Thorlabs, USA], a galvanometric mirror [OP0314024CTI, American Mirrors, USA] scanned at 400 Hz, and an illumination objective [LMPlan N 5x/0.10 IR, Olympus, Japan]. Volume scanning was achieved by moving the fiber output with a piezo actuator [PZ 400 OEM SG, Piezo Jena, Germany], thereby shifting the light sheet along the detection axis. The underlying optical design and static light-sheet characterization are described in detail in [15].

The detection unit comprised a detection objective [XLUMPLFLN20x, Olympus, Japan] mounted on a piezo positioner [NanoX 400 SG, Piezo Jena, Germany] driven by a controller [ENV 800 nanoX SG, Piezo Jena, Germany], a short-pass emission filter [FF01-750/SP-25, Semrock, USA], a tube lens [AC254-150-A, Thorlabs, USA], and an sCMOS camera [C13440 v2, Hamamatsu, Japan]. Synchronization and device control were handled with an acquisition board [PCIe-6363, NI, USA].

The mechanics for rotation comprised a rotation stage [03-0260-00-ZZ00, Rollix Defontaine, France], a servo motor [DBL3H00130-0360-000-S40, Danaher Motion, Germany], a controller [SERVOSTAR 343, Kollmorgen Seidel, Germany], and a gear reducer [NBAR075-S1-P1, Apex Dynamics, Taiwan] coupled to a rear-mounted crank-linkage mechanism. This geometry produced sinusoidal oscillation of the platform and allowed the stimulation amplitude to be adjusted mechanically. Front, back, and side views of the resulting instrument are shown in Fig. 1A.

### 4.3. Mechanical stability measurements

Mechanical stability was quantified with 1 µm fluorescent beads positioned at the sample location, using the same bead-based 3D tracking procedure described previously in [14]. Briefly, out-of-focus bead images produced concentric diffraction rings whose radial profile was used to construct a cylindrical profile over axial position. Vertical correlation of this profile across microscope positions was then used to estimate axial displacement with submicron precision, while lateral bead motion was measured from the image centroid. Bead images were acquired while the microscope was rotated through the angular range of ± 25° and repeated across time, and the resulting displacements were averaged to obtain the mean and standard deviation reported in Fig. 1C.

### 4.4 Fiber bending and polarization characterization

Bend-dependent transmission was measured with a dedicated characterization setup attached to the rotating light-sheet module (Fig. 2). The hollow-core fiber length used for characterization was 1.5 m. For bend-loss measurements, the fiber curvature was varied systematically and transmitted power was measured at the module output. For rotation-coupled polarization measurements, circular polarization was generated before fiber coupling with a quarter-wave plate, and the output was analyzed either directly or with a polarization cube while the system was subjected to sinusoidal or step rotations.

To quantify curvature-dependent polarization changes, a polarizing cube equipped with a power meter on each output arm was placed after the fiber. For each fiber configuration, the cube was rotated to minimize the signal on one arm and maximize it on the other. The cube angle provided the orientation of the output polarization state, and the residual imbalance between the two arms was used to estimate ellipticity. These measurements were used to characterize transmission stability and polarization robustness under the same dynamic conditions relevant for the microscope.

### 4.5. Vestibular stimulation and functional imaging

Larvae were mounted for light-sheet imaging and rotated together with the microscope so that vestibular stimulation was delivered physiologically about the pitch axis. For the recordings shown in Fig. 3D–F, the microscope was driven with a sinusoidal waveform of ± 10° at 0.2 Hz. Fish were paralyzed to suppress motion artefacts.

Functional imaging was performed in two-photon mode with 915 nm excitation delivered through the hollow-core fiber at an 80 MHz pulse repetition rate with precompensation configured as described for 100 fs pulse delivery in the static platform [15]. Volumes were acquired as stacks of 15 planes spaced by 10 µm. Each plane was recorded with a 29 ms exposure followed by a 1 ms delay before stepping to the next plane, corresponding to 30 ms per plane and 450 ms per volume. After each volume, an additional 50 ms delay was introduced to allow the piezo to return to the first plane. In total, 1200 volumes were recorded over 10 min, corresponding to 120 stimulation cycles.

### 4.6. Image processing and phase-map analysis

Images were corrected for lateral drift before phase-map analysis using the registration procedure described in [14]. Functional responses to sinusoidal vestibular stimulation were quantified by Fourier-based phase analysis of the fluorescence signal at the stimulation frequency, as previously described for rotating one-photon vestibular imaging [14]. Phase maps were computed at pixel resolution and represent the phase of fluorescence modulation at the stimulation frequency relative to the stimulus. The maps were visualized after anatomical registration. For multi-fish averages, individual brains were aligned to a common reference with CMTK-based registration and compared to the Z-Brain atlas [23, 24]. Average phase maps were then calculated across animals after alignment.

## Data availability

The data and analysis code supporting the findings of this study are available from the corresponding author upon reasonable request.

## Supplementary figures

**Fig. S1.**
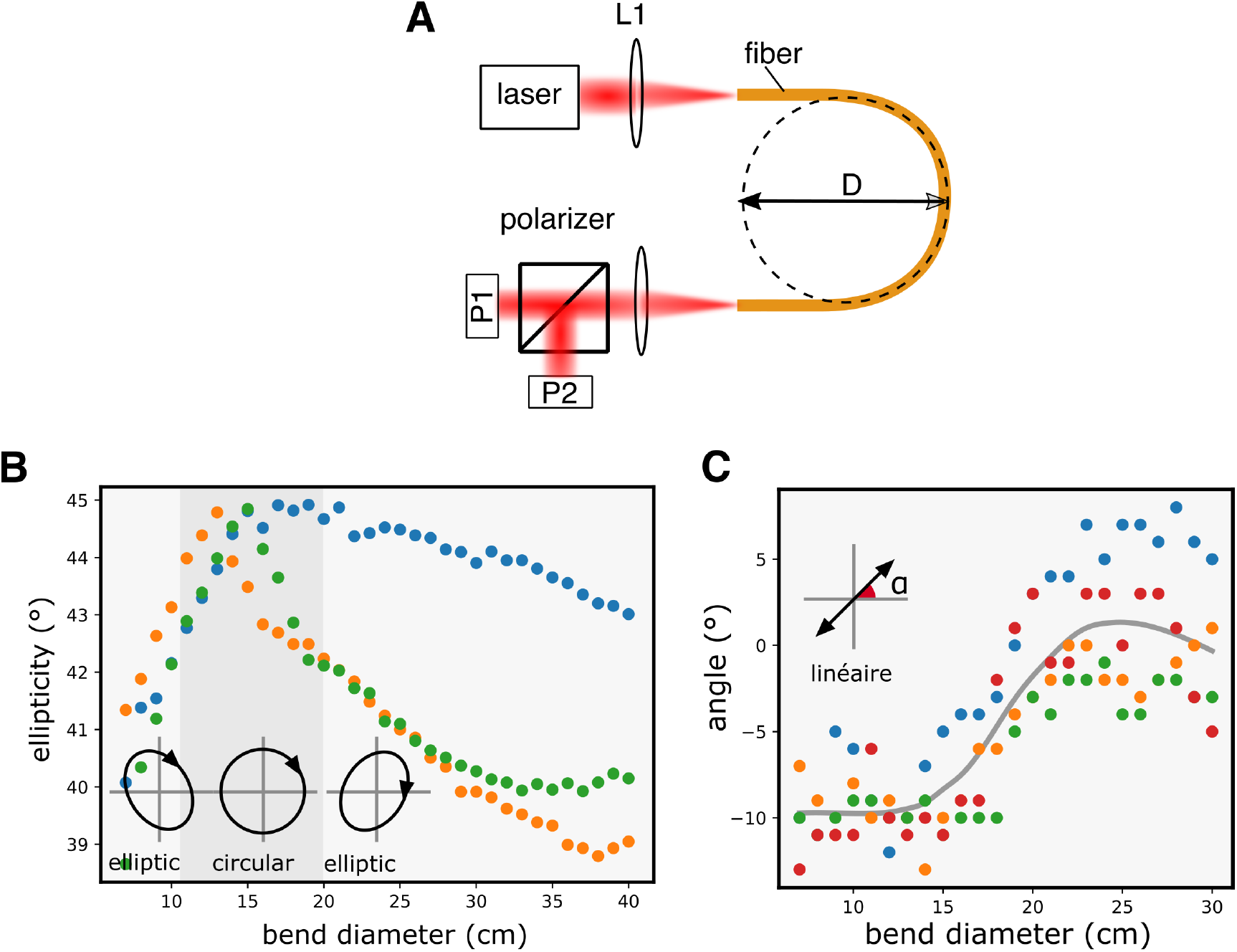
Additional polarization characterization of the hollow-core fiber under bending. (A)Measurement schemes used to quantify bend-dependent polarization changes after fiber transmission. Top: fiber-bending geometry parameterized by the bend diameter *D*. Bottom: polarization-analysis configuration in which a polarizing cube equipped with a power meter on each output arm was rotated at the fiber output to minimize one arm and maximize the other. The cube angle gave the output polarization orientation, and the residual power ratio between the two arms was used to estimate ellipticity. (B) Measured ellipticity as a function of bend diameter for repeated measurements. (C) Measured polarization angle as a function of bend diameter for the same measurements; the gray curve indicates the smoothed mean trend.

**Fig. S2.**
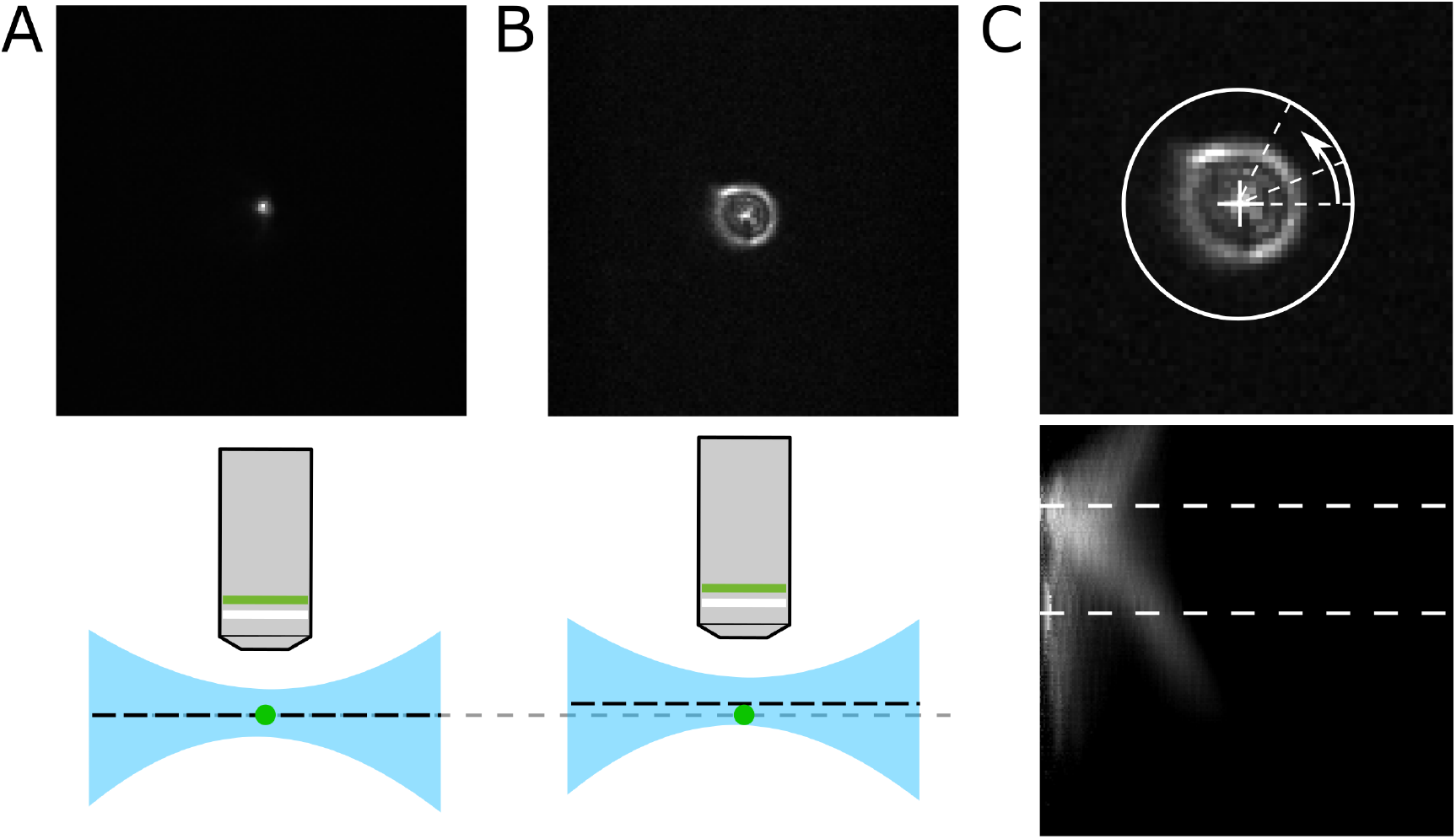
Expanded mechanical-stability measurement procedure. (A) Image of a 1 µm fluorescent bead at the sample position in focus. (B) The same bead imaged out of focus, producing concentric diffraction rings. (C) Radial averaging of the radial profile of the bead image and construction of the corresponding cylindrical profile over axial objective position. This profile was used to estimate axial displacement with submicron precision during microscope rotation, while lateral displacement was obtained from the bead centroid.

